# GlyT2-positive interneurons regulate timing and variability of information transfer in a cerebellar-behavioural loop

**DOI:** 10.1101/2024.07.10.602852

**Authors:** Ensor Rafael Palacios, Conor Houghton, Paul Chadderton

## Abstract

GlyT2-positive interneurons, Golgi and Lugaro cells, reside in the input layer of the cerebellar cortex in a key position to influence information processing. Here, we examine the contribution of GlyT2-positive interneurons to network dynamics in Crus 1 of mouse lateral cerebellar cortex during free whisking. We recorded neuronal population activity using NeuroPixels probes before and after chemogenetic downregulation of GlyT2-positive interneurons. Under resting conditions, cerebellar population activity reliably encoded whisker movements. Reductions in the activity of GlyT2-positive cells produced mild increases in neural activity which did not significantly impair these sensorimotor representations. However, reduced Golgi and Lugaro cell inhibition did increase the temporal alignment of local population network activity at the initiation of movement. These network alterations had variable impacts on behaviour, producing both increases and decreases in whisking velocity. Our results suggest that inhibition mediated by GlyT2-positive interneurons primarily governs the temporal patterning of population activity, which in turn is required to support downstream cerebellar dynamics and behavioural coordination.

**SIGNIFICANCE STATEMENT:** The cerebellum has a simple and conserved structure which has tantalised neurobiologists wishing to understand its function. Here we look at the role of granule cell layer inhibitory interneurons, Golgi and Lugaro cells, in the cerebellar cortex. We selectively turned down the activity of these cells in the awake cerebellum to characterise their influence on network activity and behaviour. We show that downregulation of Golgi and Lugaro cells has very little influence on sensorimotor representations in the cerebellum (i.e., *what* is represented), but instead modulates the timing of cortical population activity (i.e., *when* information is represented). Our results indicate that inhibitory interneurons in the granule cell layer are necessary to appropriately pace changes in cerebellar activity to match ongoing behaviour.

## INTRODUCTION

The cerebellum is implicated in a diverse range of behaviours, extending from simple reflexes to complex functions such as language, and social interaction (Baumann & Mattingley, 2022; Kelly et al., 2020; Silveri, 2021). The circuit architecture underpinning this behavioural diversity is relatively simple and conserved across its extent, suggesting that different classes of input are subject to common processing mechanisms. The cerebellar cortex lacks recurrent excitation and almost all local feedback is mediated via GlyT2-positive inhibitory interneurons situated in the granule cell layer. Golgi cells receive direct input from MFs and excitatory feedback from PFs, enabling them to exert feedforward and feedback control over granule cells (Dieudonne, 1998; Eccles et al., 1966; Kanichay & Silver, 2008). Golgi cells are numerically sparse relative to granule cells and possess extended axonal arborisations, meaning each Golgi cell innervates many hundreds or thousands of granule cells. Inhibition via Golgi cells could thus potently impact the transformation of MF signals into PF activity. *In vivo* measurements confirm that inhibition gates the transformation of sensorimotor information in the granular layer (Chadderton et al., 2004; Duguid et al., 2012), sparsifying responses to sensory input, and facilitating pattern separation at the population level (Fleming et al., 2024). Lugaro cells make up approximately one third of granule cell layer inhibitory interneurons (Simat et al., 2007), and are known to make synaptic connections onto both Golgi cells and molecular layer interneurons (Eyre & Nusser, 2016). Although comparatively little is known about their function *in vivo* (Prestori et al., 2019), their sensitivity to serotonin levels in the cerebellar cortex (Dieudonne & Dumoulin, 2000) suggests Lugaro cells could regulate information processing in the granule cell layer based on behavioural context. Currently, we lack quantitative information about how changes in Golgi/Lugaro cell activity influences cerebellar cortical dynamics and affects sensorimotor representations in the molecular and Purkinje cell layers. Indeed, the net influence of granule cell layer inhibitory interneurons is not known but may be presumed to alter both the rate and timing of granule cell output. Resultant changes in PF excitation could have net excitatory or inhibitory effects at the level of Purkinje cells, and consequently for cerebellar nuclear neurons.

Here we investigate the influence of granule cell layer inhibitory interneurons on cerebellar population activity in the context of whisker movement, to reveal how inhibitory feedback influences sensorimotor transformations and ongoing behaviour. Neuronal representations of whisking activity in the lateral cerebellum are widespread (Bosman et al., 2010; Brown & Raman, 2018; Chen et al., 2016, 2017; Zhai et al., 2024), and are partly based on a linear rate code, whereby the firing rates of cerebellar neurons predict upcoming whisker position (Chen et al., 2016). These representations are first seen in patterns of mossy fibre input and are passed forward at each stage of cerebellar cortical processing (Chen et al., 2017). Golgi cell inhibition has the potential to modulate how incoming whisking information is encoded and transformed within the granule cell layer (Gurnani & Silver, 2021; Palacios et al., 2021). However, it is unknown whether, and how, changing granule cell layer inhibition impacts neuronal representations of whisker behaviour, and indeed whisking behaviour itself. We have recorded whisker movements and population activity in the lateral cerebellum using NeuroPixels probes in head-fixed mice expressing inhibitory DREADDs (designer receptors exclusively activated by designer drugs) selectively in Golgi and Lugaro cells (i.e. GlyT2-positive neurons) of lobule Crus 1. In these mice, it was possible to downregulate GlyT2-positive cell activity by activating DREADDs with the agonist, clozapine N-oxide (CNO), applied topically on the recording site. Our results reveal a subtle but direct influence of GlyT2-positive cells on network dynamics and consequently, the cerebellar-behavioural loop underlying voluntary whisking.

## MATERIALS AND METHODS

Experiments were conducted on male and female C57BL/6 (wild-type) and heterozygous C57BL/6-Slc6a5<tm1.1(cre)Ksak> (GlyT2-Cre knockin) mice (Kakizaki et al., 2017) aged between 3-6 months. Animals were housed under a 12/12 h reversed light–dark cycle with food and water available *ad libitum*. Habituation and recordings were performed during the dark phase of the cycle. The C57BL/6-Slc6a5<tm1.1(cre)Ksak> mouse strain was provided by the RIKEN BRC through the National BioResource Project of the MEXT/AMED, Japan. All experiments were conducted under the United Kingdom Animals (Scientific Procedures) Act 1986 in accordance with Home Office guidelines.

### Surgical procedures

Mice were anaesthetised for all surgical procedures. Mice were placed in an induction chamber, into which 5% v/v isoflurane (Harvard Apparatus Ltd) was administered via inhalation using oxygen (1.2–1.6 L/min). Once anaesthetized the concentration of isoflurane was lowered to 1–2% v/v to maintain anaesthesia for the duration of the surgery. The body temperature of the mouse was maintained at 37 ± 0.5 °C using a homeothermic heat mat (DC Temperature Control System, FHC). Corneal drying was prevented using an ocular lubricant (Lacrilube, Allergen). To induce analgesia, mice were injected subcutaneously with carprofen (Rimadyl, 5 mg/kg), buprenorphine (Vetergesic, 0.1 mg/kg) and lidocaine (2 mg/kg, locally). Surgery was carried out in a stereotaxic frame under aseptic conditions to prevent infections. After surgery, mice were placed in a warmed box (∼37°C) and allowed to recover for as long as required. Mice were then returned to their home cages and monitored closely for at least 24 hrs.

GlyT2-Cre mice underwent two surgical operations: one to transduce cre-dependent expression of DREADDs in the cerebellar cortex via viral injection, and one to prepare the animal for head-fixed extracellular recording. During the first surgical session, a small craniotomy (∼1 mm diameter) was performed over Crus 1 (6.36 mm posterior and 2.5 mm lateral of Bregma). AAV9-hSyn-DIO-hM4D(Gi)-mCherry (500-1000 nl) was injected with a glass pipette at different depths (starting at 600μm retracting to surface in steps of 100μm), waiting 2 minutes between pipette retractions. At the end of the operation, skin over the head was sutured and animal was allowed to recover.

The second surgical session took place 8 weeks later to enable ample expression of the DREADD receptor protein. First, neck muscles were gently moved to uncover the cerebellum. A custom-made head implant was fixed to the exposed cranium using tissue glue (Histoacryl, Braun Corporation) and dental cement (Associate Dental Products Ltd). A small craniotomy (∼1.5 mm diameter) was performed over the right cerebellar hemisphere, the dura was removed, and a reference screw was inserted in contact with the underlying brain. To secure both head place and reference screw, dental cement (Associate Dental Products Ltd) was used to cover the skull, sparing a recording well above Crus 1 of the left cerebellar hemisphere. A second craniotomy (∼2 mm diameter) was performed over left Crus 1, and the dura was removed. Finally, a layer of agarose (1.5% in phosphate buffered saline; PBS) was used to cover the brain, a layer of Kwik-Seal was used to protect the brain, and a layer of nail polish was used to fix the Kwik-Seal. Wild-type mice underwent only the second surgical operation in preparation for recordings. All mice had their whiskers trimmed on the left (recording) side, preserving whiskers C1, C2 and C3 (posterior whiskers on the third row from top).

### Electrophysiological and video recordings

Mice were head-fixed in the recording apparatus in preparation for electrophysiological recording. The Kwik-Seal was removed, the surface of the brain was cleaned from agarose and kept moist with PBS. NeuroPixels probes were fixed to a micromanipulator (IVM, Scientifica) via a custom-made 3D printed holder. Each probe was coated with DiI stain (2.5 mg/ml) for *ex vivo* probe tracking before insertion at an angle perpendicular to the cortical surface. The probe was advanced into the cortex at a speed of 2-5 μm/s to a depth of 2500-3000 μm. Thereafter the probe was retracted for ∼100 μm and left to settle for 10-15 min, allowing the brain to relax. Under infrared light illumination, whisker movements were filmed with a high-speed camera (Genie HM640; Teledyne Dalsa Inc, USA) operating at 299 frames per second. Video acquisitions were controlled by Streampix 6 software (Norpix, Canada). The open-source software spikeGLX 3.0 (https://billkarsh.github.io/SpikeGLX/) was used to record NeuroPixels data at 30 kHz. Video and electrophysiological recording were synchronised by a TTL pulse originating from the video software. In chemogenetic experiments, a baseline of 10 or 20 min was recorded prior to topical administration of CNO (30 μM), delivered to the recording well. Previously Stachniak et al., 2014 showed that intracranial injection of 1 μM CNO is already sufficient to reliably inhibit presynaptic neurotransmitter release within minutes. After CNO or vehicle delivery, electrophysiological recordings continued for a further 40-50 mins, for a total recording time of 1 hour. Animals underwent two recording sessions, the first performed at least 4 hours after recovery from the craniotomy, the second on the following day. After recording, the probe was extracted, rinsed with deionised water and left in freshly made 1% tergazyme solution for at least 24 hours. After the first recording, the exposed brain surface was first rinsed with PBS, then covered with agarose, and finally protected with a layer of Kwik-Seal that was fixed to the surrounding dental cement with nail polish. Mice were returned to their cage. After the second recording session, mice were humanely killed. See **Table 1** for full information on data included for analysis.

### Electrophysiological data processing

Output files (.imec) were preprocessed using the command-line tool CatGT (https://billkarsh.github.io/ SpikeGLX/#catgt) to apply a high pass filter (cut-off 300 Hz) and global demux filters, a common average referencing that takes in account the probe channels subgrouping during data acquisition. Spike sorting was conducted using the open-source software Kilosort 2 (https://github.com/MouseLand/ Kilosort) to group spikes into units, each representing the activity of a distinct putative neuron. Manual curation of sorted units was done with the open-source Python-library Phy2 (https://github.com/cortex-lab/phy), and consisted in merging and splitting units, as well as categorising them into ‘good’, ‘bad’ and ‘mua’ (multiunit activity) units. In all analysis, only good, i.e. well-isolated, units located in the cerebellar cortex were used.

### Video processing and behavioural analysis

Video output (.avi) files were transcoded (.mp4, libx265 encoder), re-sampled at 299 Hz) and cropped using the ffmpeg software. We used the open-source toolbox DeepLabCut (https://github.com/DeepLabCut/DeepLabCut) to label four landmarks on whiskers C1, C2 and C3 (**Figure 1a**) across the entire recording; these landmarks were then used to compute whisker position. Specifically, the arctan function was used to measure the azimuthal angle between the first whisker segment (traced from the first, basal marker to the second marker) and the horizontal line passing through the basal marker. Increases in angle correspond to whisker protraction, whereas decreases correspond to whisker retraction. Next, the phase of the whisking cycles was extracted using the Hilbert transform, in order to compute the slowly changing whisking amplitude and setpoint across cycles (Hill et al., 2011). Finally, the whisking amplitude was low pass filtered and used to identify periods of whisking. A heaviside function with a threshold of 10°, was used to discriminate whisking from resting periods. All whisking analysis focused on the C3 (most anterior) whisker.

**Figure 1.**
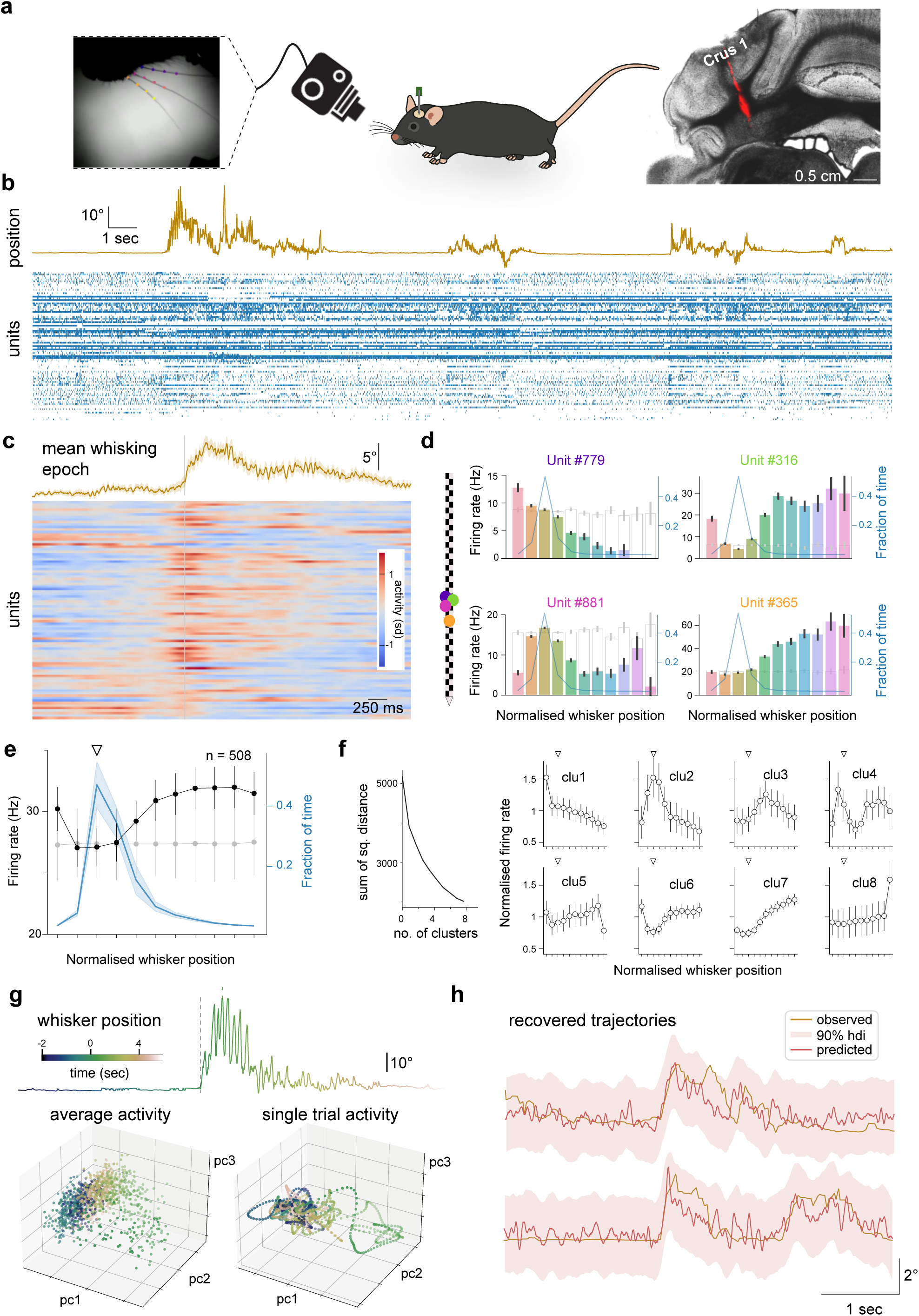
Single unit and population activity in Crus 1 during voluntary whisking. **a.** Monitoring whisker movement and neuronal population activity in lateral cerebellar cortex using high-speed videography and NeuroPixels recording in head-fixed mice. Left: Whiskers were labelled and reconstructed using DeepLabCut to recover movement trajectories. Image credit: Elisabeth Meyer. Right: Brightfield image of coronal cerebellar section superimposed fluorescent image tract left by NeuroPixels probe coated with DiI localised to lobule Crus 1. **b.** 30 second segment of simultaneous whisking (top) and population activity (bottom; *n* = 88 units) from a single recording session. **c.** Trial-averaged whisker position and neuronal activity from the same recording, aligned to onset of whisking bout. **d.** Tuning curves of four individual cerebellar units from the same recording. Each unit displays a different relationship between firing rate and whisker position. Blue lines indicate fraction of time spent at different whisker positions; empty grey bars show tuning curves for shuffled data. Left inset: colour-coded centre of mass of each unit on NeuroPixels probe. **e.** Average tuning curve for all recorded units, from all recordings (black line; n = 508, N = 12). Population firing rate increases monotonically from whisker resting point (white triangle) in both protractive and retractive directions. Blue line indicates fraction of time spent at different whisker positions; shading represents standard deviation; white triangle indicates whisker resting point; grey line shows tuning for shuffled data. **f.** Average tuning curves for all units clustered in eight groups. Units were clustered using k-mean algorithm based on individual unit tuning curves. Heterogeneity in tuning curves enables a continuous encoding of whisking position. **g.** Single trial of whisker movement (top) with corresponding projections in 3D principal component space for average (bottom left) and single trial (bottom right) population activity. Population activity in this space has structure that reflects whisking dynamics, both on average and at a single trial level. **h.** Recovered movement trajectory using cerebellar population activity. Reconstruction of the whisking setpoint using a linear combination of the first three principal components computed from neuronal populations. Whisker setpoint (brown line) during two trials, together with predictions (red) and the highest density interval (hdi, red shading).

### Histology

Brains were extracted and left in 4% PFA solution for 24-48 hours at 4°C. After washing with PBS, cerebellar coronal sections (50-100 μm) were sliced using a vibrating microtome (Leica VT1000S). Fluorescent images were acquired using a confocal microscope (Leica DM4000 B).

## EXPERIMENTAL DESIGN AND STATISTICAL ANALYSES

### Single unit analysis

To compute the tuning curve for each unit, we computed the total spike count and average whisker position (angle) in time bins of 33 ms. We then discretised the average whisking angles into 11 bins, ranging from the minimum to the maximum angle observed in each recording and paired the spike count in each time bin with the corresponding whisker angle. These data were finally used to plot the firing rate, together with their standard error, across the 11 angle bins.

Both the entropy of and KL-divergence between the pre- and post-drug tuning curves for each unit was computed using the function entropy from the scipy.stats library. The tuning curve for each unit was normalised to be the probability mass function of firing rates over whisking position.

Clustering of tuning curves was performed with *k*-means algorithm (sklearn.cluster.KMeans package), used on the spline coefficient fitted to the individual unit tuning curves. The estimation of the tuning curves can be noisy due to, for example, errors during the spike sorting process, which may lead to the contamination of the spike history assigned to one unit by the activity of nearby neurons. This noise, in turn, could affect clustering of tuning curves into meaningful groups. To reduce the impact of this source of noise on our clustering analysis, we fitted a spline model to each tuning curve: the model used seven cubic b-spline functions to smooth the firing rate associated to each angle bin based on the firing rate of adjacent bins, while maintaining the underlying structure of the tuning curves. The smoothed tuning curves were then clustered using a *k*-means algorithm with *k* = 8. We finally computed the average tuning curve from units within each cluster. Results were qualitatively the same with fewer than *k* = 8 clusters.

### Principal component analysis of population activity

We used principal component analysis to describe the population activity of each recording (number of units per analysis can be found in **Table 1**). Projections onto the first three eigenvectors (principal components; pcs) described the first three orthogonal axes in neuronal space along which population activity varied the most. To perform this analysis, we first segmented each recording into discrete whisking epochs (from -2 to +3 seconds around the start of each whisking bout, defined using behavioural analysis above), and computed the trial-averaged firing rate for each unit. Firing rates were used to compute correlation matrices across units, from which eigenvalues and eigenvectors were derived. Eigenvalues indicate the amount of neuronal activity variability captured along the axis specified by the corresponding eigenvectors. For each recording, the eigenvectors had length equal to the number of units present in that recording. The values of an eigenvector are known as loads and correspond to the weight associated to each unit when linearly combining the firing rate from all units; hence, the loadings of an eigenvector correspond to how much each unit contributes to the pc associated with that eigenvector. The eigenvectors can be used to project either each unit’s firing rates in pc space or single trial activity: to compute the pc for single trial activity, we first smoothed single trial spike counts with a Gaussian window of std = 20 (we also used std = 5 but results were equivalent; time bin width = 3.3 ms).

Cross-correlation between whisking position and each pc was obtained using the scipy.signal.correlate package. All cross-correlations were normalised by the absolute maximum value of the first pc. For analysis involving CNO manipulation, cross-correlation peaks for pre- and post-drop periods were computed, and the respective distributions were compared using a two-sided Wilcoxon signed-rank test. To compared cross-correlations across conditions, we first computed the within-recording contrast between post-versus pre-drop cross correlation peaks; next, for each pc and experimental condition, we fitted the contrasts with a Gaussian distribution, and finally we took the difference in posterior samples of the mean between GlyT2-CNO and control conditions.

### Analysis to assess efficacy of DREADD manipulation

A multilevel model was used to account for the variability in total cerebellar cortical spike counts detected over time across recordings; counts were the sum of the spikes assigned to good (i.e. well-isolated) cortical units. Spike counts were computed for successive 5 minute-time bins, with the 0-time bin including the time of drug application. The counts in each bin were scaled with respect to the count in the -5- minute bin.

In the model, spike counts were described using an inverse-Gamma distribution:

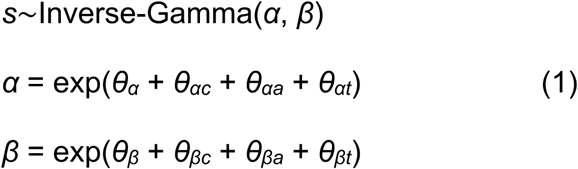

The inverse-Gamma distribution is used because it provides a good fit to the population spike count data (**Figure 3e**); the two parameters *α* and *β* are usually referred to as the shape and scale parameters and are related to the mean and variance by *μ* = *β*/(*α* − 1) and *σ*^2^ = *β*^2^/[(*α*−2)^2^(*α*−1)]. Here *α* and *β* are each linked to a linear model by a loglink function. Both linear models involve a coefficient *θ* for each explanatory variable, namely, the experimental condition *c*, the time of the recording *t* (dim = 9), and the animal ID *a* (dim = 17). The experimental conditions were, (i) application of CNO to cerebellar surface of DREADD-transduced GlyT2-Cre mice (GlyT2-CNO, *n* = 19 recordings), (ii) application of CNO to cerebellar surface of wild-type mice (WT-CNO, *n* = 5 recordings), (ii) application of PBS the cerebellar surface of wild-type or GlyT2 mice (WT-Veh, *n* = 9 recordings). Experimental conditions were split by recording period into a pre-drug or baseline period (bins before 0-time bin) a post-drug period (0-time bin and onward), for a total dimensionality of 3 x 2 (dim=6) for the *θ_αc_* and *θ_βc_* coefficient. The contrasts between pre- and post-drug coefficient *θ_c_* were used to assess the effect of the experimental manipulation.

**Figure 2.**
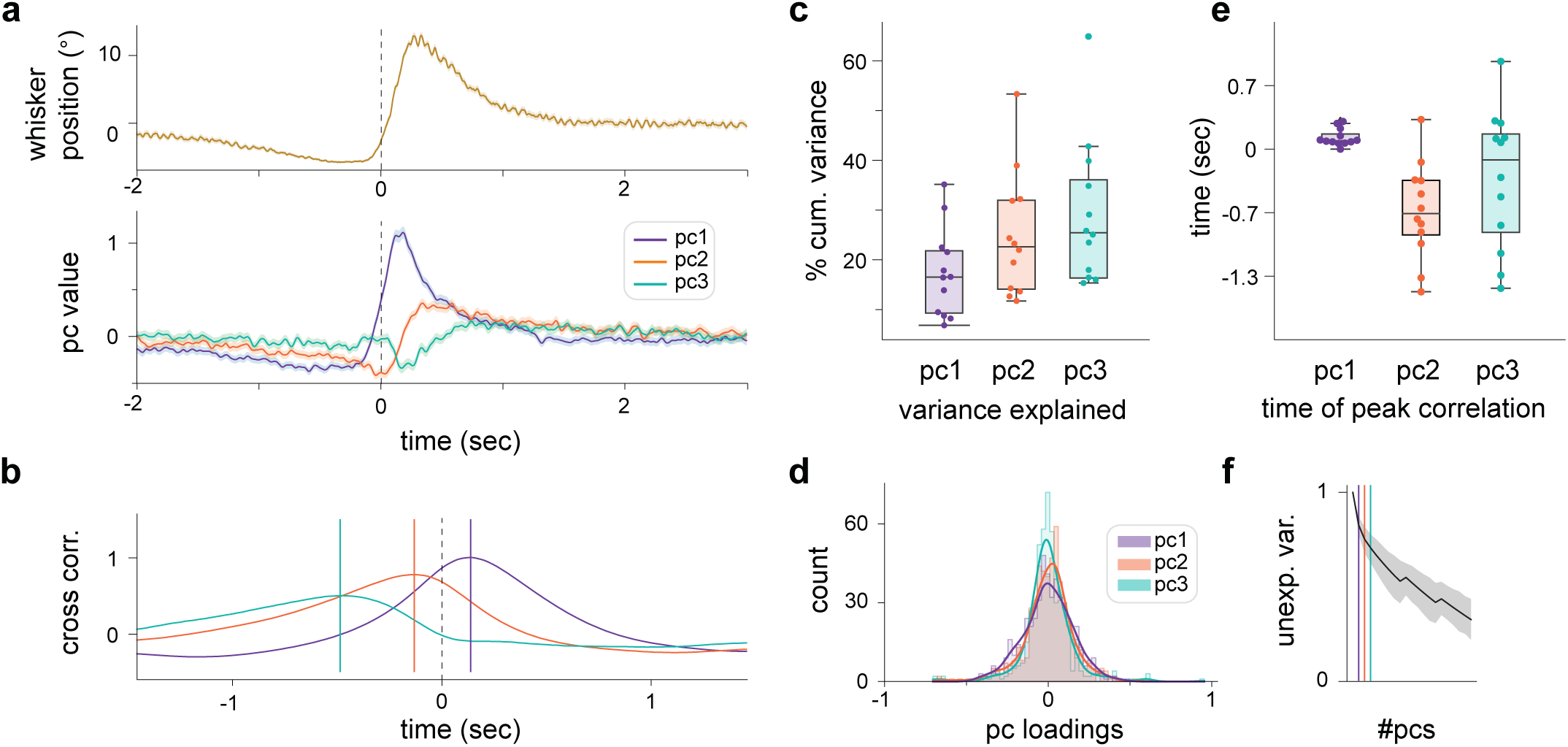
Population representations of whisker movement at different timescales. **a.** Trial-averaged whisking position (top) and projected population activity (bottom) for one recording. **b.** Cross-correlations between pc1-3 and whisking activity for the same recording; vertical lines indicate times of peak correlation. **c.** Cumulative variance explained by pc1-3 across all recordings (*N* = 12). Only a modest amount of variability in neuronal activity is accounted for by first three principal components. **d.** Distribution of unit loadings (*n* = 508) for pc1-3. Kurtosis measurement, to quantify the number of outliers in each distribution, reveals fewer outliers for pc1 than what would be expected if the data were normally distributed (excess kurtosis -2.76). This indicates that information contained in pc1, which best reflects whisking activity, tends to be distributed across neurons. The loading distributions for pc2 and pc3 have respectively a similar or higher number of outliers compared to normally distributed data (excess kurtosis -0.43 and 4.97, respectively), indicating that information in pc2 and pc3 is increasingly concentrated in fewer units. **e.** Time of peak correlation for pc1-3 across all recordings. The distribution for pc1 is tightly concentrated around ∼40 ms, meaning that information captured by pc1 tends to anticipate whisking activity with high temporal precision (one sample *t* test, *t* = 4.71, *p* = 0.0006). The information contained in pc2 and pc3 shows the opposite trend, with more variability (pc2: mean -194 ms, *t* = -4.44, *p* = 0.0009; pc3: mean -92 ms, *t* = -1.44, *p* = 0.17), suggesting that these components might reflect different aspects of behaviour. **f.** Mean decrease in unexplained variance (unexp. var.) across recordings with increasing numbers of principal components (#pcs); standard deviation shaded in grey.

**Figure 3.**
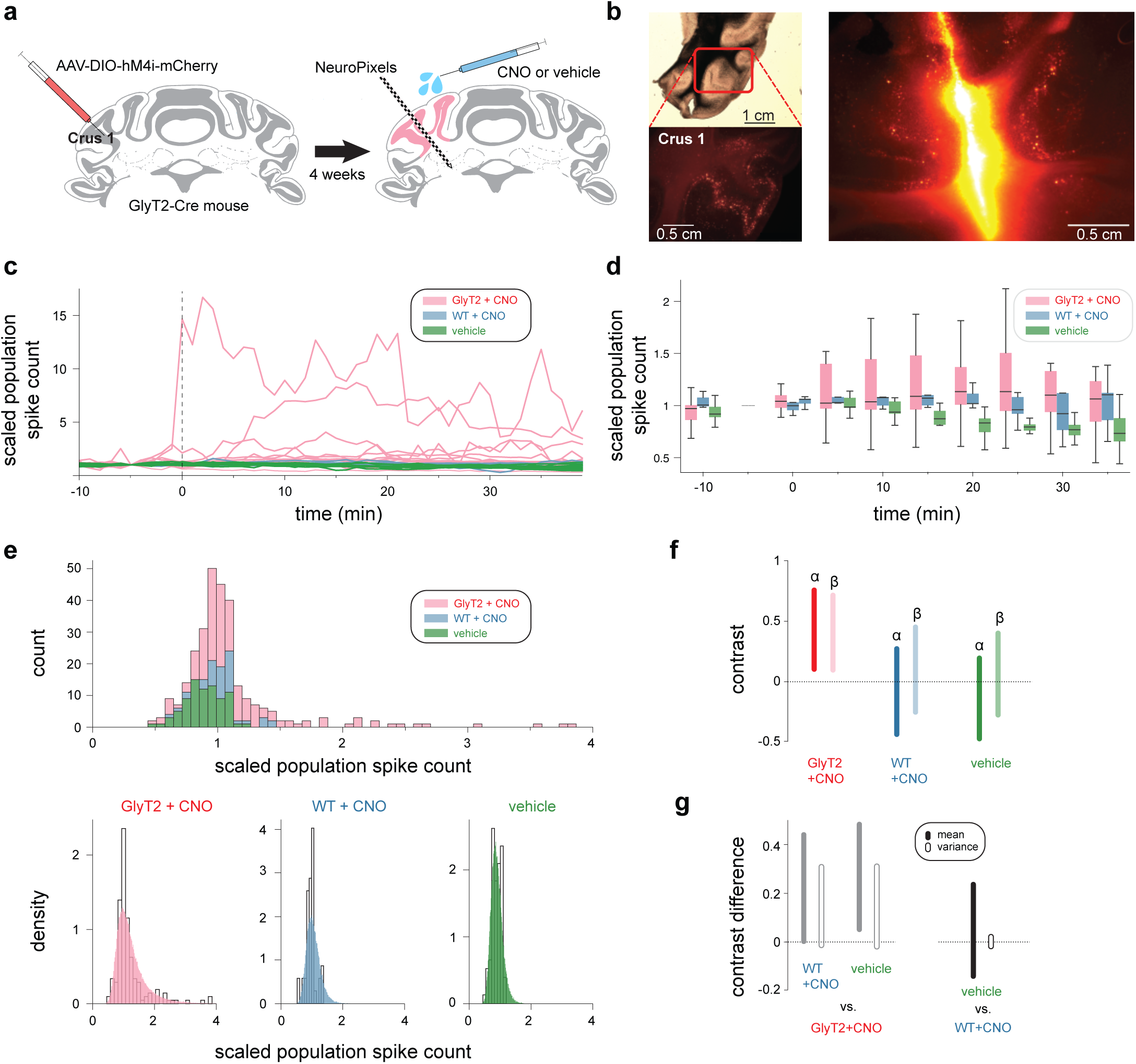
Chemogenetic inhibition of GlyT2-positive cells increases neuronal activity in cerebellar cortex. **a.** Left: targeted expression of hM4Di receptors was achieved via injection of AAV-DIO-hM4Di-mCherry into the lateral cerebellar cortex of GlyT2-Cre mice, selectively expressing Cre-recombinase in GlyT2-positive cells in this brain region. Right: NeuroPixels probes were targeted to site of viral injection. During electrophysiological recordings, the exogenous drug, clozapine-N-oxide (CNO) was topically applied to activate hM4Di receptors. **b.** Left: GlyT2-Cre mouse brain selectively expressing hM4Di in GlyT2-positive cells of the lateral cerebellar cortex. Right: tract left by NeuroPixels probe coated with DiI, showing co-localisation of the site of recording and hM4Di expression. **c.** Changes in population spike count before and after CNO/vehicle delivery (at 0 min) normalised by count at -5 min, for three experiment conditions. Pink, GlyT2-Cre mice and CNO (*N* = 19); Blue, C57BL6 mice and CNO (*N* = 5); Green, GlyT2-Cre/C57BL6 mice and saline vehicle only (*N* = 9). **d.** Box plots showing pooled data for each experimental condition. **e.** Multilevel modelling approach to assess effect of drug delivery on cerebellar population activity. Population spike count distributions for each condition (top) were modelled using an inverse-Gamma distribution, described by shape parameter α and scale parameter β. Each parameter was modelled as a linear combination of different coefficients, including two θ_cond_, one for α and one for β, which captured the specific effect of the experimental manipulation on total spike counts. Comparing the empirical distribution with the distribution of posterior predictive checks (samples from the model after fitting) shows that the model captures the overall structure of the data (bottom). **f.** 94% highest density intervals (hdi) of the contrasts between post- and pre-drop posterior samples for θ_cond_ for α and β. The contrasts highlight a specific effect of GlyT2-positive cell manipulation (GlyT2 + CNO) on both α and β parameters. **g.** Contrast difference (mean and variance) between post- and pre-drop of the inverse-Gamma distribution fitted to total spike counts. Bars indicate the 94% hdi of the posterior differences. Left, contrast difference between GlyT2 + CNO condition and each control condition. Right, contrast difference between the two control conditions. GlyT2-positive cell manipulation produces positive contrast differences versus both control conditions.

The additional explanatory variables, *t* and *a*, were used to explain respectively spike count variability within each pre-drug time bin due to natural changes in network activity over recording time (*n* = 9, excluding -5-min time bin), and variability due to the particular conditions of the network in each animal (*n* = 17), which might be affected, for example, by the surgical procedure. Thus, the addition of these explanatory variables allowed us to test within conditions for a difference in pre- and post-drop spike count distribution, after accounting for variability explained by temporal correlations and between-animal differences.

Each *θ*, except *θ_t_* was drawn from a vector of independent normal distributions, of length equal to the dimensionality of *θ*. The prior distribution for *θ_t_* was a multivariate normal distribution with covariance decaying with time distance, accounting for temporal correlations within the data:

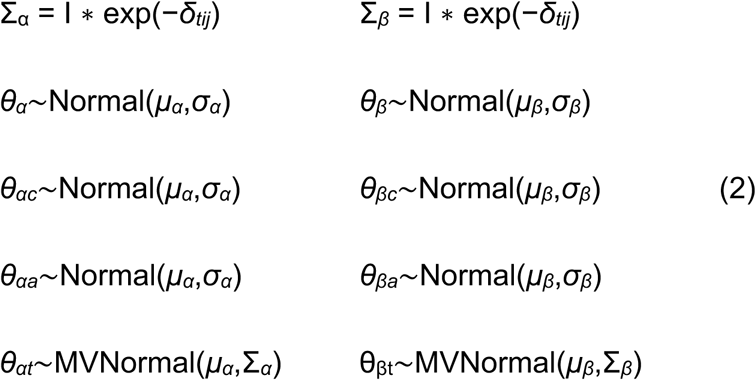

with *δ_tij_* between any two periods *i* and *j* is defined as:

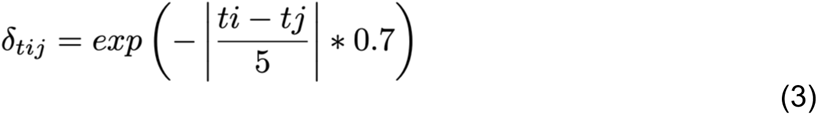

Where|*t_i_* – *t_j_*|/5is the absolute time difference between period *i* and *j* scaled to unity (5 is the time width of each period in minutes).

Finally, the hyperpriors:

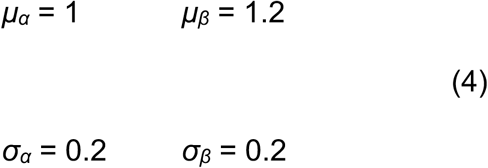

were chosen to sample prior predictive samples within a similar scale of the observed population spike counts, including extreme values.

The posterior samples for the mean and variance spike count parameters were used to compute the difference across conditions between post- and pre-drug contrasts in mean and variance due to the experimental manipulation (**Figure 3f**). In practice, the contrasts were computed using the same posterior samples, except for *θ_αc_* and *θ_βc_*, which differed for pre- and post-drug mean and variance. The difference between contrasts was used to assess changes in mean and variance across conditions (**Figure 3g**).

### Analysis of DREADD manipulation on neuronal and whisking behaviour

Comparisons were made from the pre- and post-drug time periods. The pre-drug period lasted 20 minutes all but five recordings, where this period was 10 minutes. The post-drug time period matched the pre-drug period and started 5 minutes after drug application. Recordings that exhibited little or no whisking activity were excluded from this analysis leaving N = 25 recordings in total.

For each population recording, we computed the absolute peak of the trial-averaged neuronal activity (PETH) aligned to whisking onset. The subwindow used to compute the absolute peaks spanned -0.7 to 1.3 sec around whisking onset, to focus the subsequent analysis around the initial whisking protraction period. We used the absolute peak of the PETH as a proxy of the timing of the neuronal response to whisking-related input. For each recording we computed the standard deviation (std) of the peaks, for pre- and post-drug data. For each experimental condition, the pre- and post-drug stds were compared using a two-sided Wilcoxon signed-rank test.

For corresponding whisking data, we used a linear model to fit a line to the average whisking position pre- and post-drug. Whisking data were delimited by a time window ranging from -0.06 to 0.21 sec centred around whisking onset; the time window was chosen to focus on the period of initial whisking protraction. We computed the difference between post-drop and pre-drug slopes (i.e. velocity to compare the effects of Golgi cell manipulation on the whisking protraction phase. To test whether the variance of the distribution slope contrasts in the two experimental conditions were different, we used Levene’s test (scipy.stats.levene).

### Code accessibility

Code used for the analysis is freely available here: https://zenodo.org/doi/10.5281/zenodo.13327395

## RESULTS

### Population dynamics in lateral cerebellum during voluntary whisking

To understand the contribution of GlyT2-positive interneurons to cerebellar cortical dynamics, we first set out to describe how cerebellar populations represent sensorimotor information while Golgi and Lugaro cell inhibition was intact. We recorded cerebellar population activity using NeuroPixels probes in lobule Crus 1 of awake mice and performed spike sorting to isolate single units (see Materials and Methods). We isolated an average of 42.3 (std = 25.3) cortical units per recording (*N* = 12), for a total (*n*) of 508 putative units. To correlate spiking activity with whisker movement, we simultaneously tracked spontaneous whisking via a high-speed camera (**Figure 1a**). In all cases, bouts of whisking activity were associated with pronounced alterations in cerebellar spiking activity (**Figure 1b, c**). Single cell recordings previously revealed widespread tuning to whisker position in granule cells, inhibitory interneurons and Purkinje cells (Bosman et al., 2010; Brown & Raman, 2018; Chen et al., 2016, 2017; Zhai et al., 2024). We therefore measured tuning to whisker position amongst single units of our population recordings. On a unit-by-unit basis, we examined the relationship between firing rate and whisker position, to construct individual tuning curves (see Materials and Methods). Consistent with previous work, we observed robust tuning to whisker position, such that single units exhibited their highest firing rates at preferred angles of whisker deflection.

Within locally recorded populations, we observed substantial diversity in the tuning to whisker position. Indeed, units recorded in the same penetration displayed a broad range of preferred angles and tuning profiles, including monotonic increases and decreases to whisker protraction/retraction (**Figure 1d)**. We then recombined all single unit spiking responses to calculate population tuning. Whereas individual units exhibited diverse selectivity, the mean tuning function for the population displayed a near-monotonic dependence upon whisker position, with rate changes in proportion to the magnitude of protraction and retraction (**Figure 1e**).

To further assess tuning diversity at the population level, we clustered units based on their tuning curves, testing whether we could discriminate discrete functional subclasses of unit. We fitted a spline model with 7 cubic b-spline to each tuning curve (*n* = 508, *N* = 12) to capture tuning curve shape and reduce variability due to noise (see Materials and Methods). We then inspected the output of *k*-means clustering with different numbers of expected clusters. With increasing cluster number, there was no evidence of a change in the sum-of-squared-distances decaying pattern, or elbow, which can be used as an indicator of the optimal number of clusters to be considered. Visual inspection of the mean tuning curves for each cluster revealed a progressive shift of preferred position from the most retracted to the most protracted whisker locations (**Figure 1f**). Projection of tuning curves in 3D space using t-SNE, a non-linear dimensionality reduction technique particularly suited to uncovering clustering structure further confirmed that different groups of clusters are not well isolated (data not shown). Thus, our analysis does not support discrete classes of tuning curve but rather confirms that cerebellar neurons form a functional continuum in the space of possible tuning curves. At the local population level, Crus 1 cerebellar cortical neurons heterogeneously encode whisking position, but represent increasing protractive and retractive movements via accumulating increases in firing rates.

Having established that concerted population activity is required for accurate sensorimotor representations, we performed principal component analysis upon the grouped spike trains from each recording. We initially preserved projections of population activity onto the first three eigenvectors, which describe the three orthogonal axes in neuronal space along which activity varies the most (**Figure 1g**). Analysis was restricted around whisking bouts (-2 to +3 seconds about whisking onset). Projections of population activity revealed clear structure in neuronal space during transitions from quiescence to whisking activity and back again. This structure in turn suggested that population activity contains accurate information about whisking dynamics. We tested this proposal by linearly combining the first three principal components, attempting to reconstruct movement trajectories (see Materials and Methods). Using this approach, it was possible to decode whisking setpoint during single trials (**Figure 1h**). In summary, our results show that the population activity in the lateral cerebellar cortex accurately reflects upcoming slow whisking dynamics, even when approximated within a low-dimensional space.

To quantify the relationship between different principal components and whisking, we computed the normalised cross-correlation between whisker angle and each of the first 3 principal components (pc) in each recording. Interestingly, pc1-3 showed distinct temporal relationships with respect to whisker movement (**Figure 2a,b**). Information related to whisker position contained in pc1 tended to anticipate behaviour, whereas information in pc2 and pc3 lagged behaviour. Across all recordings (*N* = 12), the peak signal of pc1 always preceded the whisking signal, indicating that, across recordings, variation in the whisking-aligned average neuronal activity expressed by the pc1 tends to anticipate behaviour (one sample *t* test, *t* = 4.72, *p* = 0.0006, mean = 36 ± 8 ms). On the other hand, the peak signal for pc2 almost always occurred after whisker movement, indicating that whisker-related information contained in pc2 lags whisking activity (*t* = -4.43, *p* = 0.001, mean = -195 ± 44 ms). Finally, peak signals for pc3 were broadly distributed around 0 ms (*t* = -1.38, *p* = 0.19, mean = -89 ± 65 ms). Notably, pc1 peaks across recordings are narrowly distributed in time; this suggests that pc1 specifically contains precise information about whisking behaviour, i.e. upcoming movement, occurring ∼40 ms in the future (**Figure 2c**).

Whisking-related information contained in pc1-3 could originate from a small proportion of units in the population, whose activity may be particularly responsive to whisking; alternatively, information could be more distributed, whereby all units contribute to some extent to the population encoding of behaviour. To measure how behavioural information was distributed across the population, we measured the pc loading of each unit (*n* = 508), indicating the contribution of a unit to the corresponding pc (see Materials and Methods). The loading distribution for each pc was centred around 0 and was roughly symmetric, meaning that units could contribute both positively or negatively to each pc (**Figure 2d**). The number of outliers in each distribution was used to quantify the extent to which each pc is dominated by the activity of few units. We calculated the excess kurtosis of each distribution, which is a measure of the frequency of outliers observed, using as a point of reference the frequency expected from normally distributed data. The distribution for pc1 had an excess kurtosis of -2.85, indicating infrequent outliers; this in turn suggests that the contribution of units across recordings to pc1 is broadly distributed within the population. In contrast, the excess kurtosis for pc3 was 4.54, indicating a high frequency of outliers, and thus a subset of units which tend alone to contribute mostly to pc3. The excess kurtosis for pc2 was -0.47, meaning that outliers are neither frequent nor infrequent. Together, these results suggest that for pc1, and thus accurate representation of whisking behaviour, information is broadly distributed across the entire population. Finally, we noted that the first three eigenvalues accounted for only a moderate amount of the total variance in the population activity (∼30%) across all recordings (**Figure 2e,f**). Therefore pc1-3 provide only a coarse description of population activity, and whisker movement only partially describes neuronal dynamics in Crus 1 despite restricting our analysis to the initial period of whisking activity. Indeed, the large proportion of neuronal variability not captured in the first 3 principal components supports the proposal that cerebellar cortex may represent additional behaviourally relevant variables.

### Chemogenetic downregulation of GlyT2-positive interneurons elevates population activity

Having established the relationship between population activity and slow whisking dynamics under baseline conditions, we next sought to understand how Golgi and Lugaro cell inhibition contributes to these representations in the cerebellar cortex. We adopted a chemogenetic strategy to selectively downregulate GlyT2-positive neuronal activity in an otherwise intact network while simultaneously recording population activity and whisker movement. Manipulation of GlyT2-positive cells was achieved via activation of the inhibitory DREADD, hM4Di, which causes reductions in excitability and neurotransmitter release (Stachniak et al., 2014) in cells expressing these receptors. Localised expression of DREADDs (**Figure 3b**) was achieved by injecting the AAV-DIO-hM4D(Gi)-mCherry virus into Crus 1 of GlyT2-Cre mice (cre-recombinase restricted to GlyT2-positive cells in the cerebellar cortex; see Materials and Methods; Kakizaki et al., 2017). DREADDs were activated by local, topical delivery of the agonist, clozapine-N-oxide (CNO), on top of the Neuropixels penetration site, which itself had been targeted as the site of virus injection (**Figure 3a**); this method was used to minimise potential off-site effects of the agonist and afforded within-minute temporal precision to our manipulation. To confirm the fidelity of our approach, we first tested the effects of topical delivery of muscimol (1.7 μg/μl), a GABA_A_-receptor agonist, during NeuroPixels recordings from wildtype mice (N = 3). In all cases, within minutes of drug application, the total spike count recorded across the cerebellar cortex started to decrease and reached its minimum within 10-30 minutes (data not shown). These results supported the viability of applying CNO topically onto cerebellar cortex to locally activate DREADD receptors.

We next examined whether CNO-dependent reduction in GlyT2-positive cell activity had a measurable impact on cerebellar cortical population activity. To do this, we compared changes in cortical spike count before and after CNO administration across three experimental groups. GlyT2-Cre mice, pre-injected with AAV9-hSyn-DIO-hM4D(Gi)-mCherry virus into Crus 1, or uninjected C57BL6 mice received topical delivery of CNO (GlyT2-CNO; *N* = 19 recordings) or saline vehicle (Veh; *N* = 9 recordings) to the cerebellar cortex. A third group of uninjected wildtype (C57BL6) mice received topical delivery of CNO to the cerebellum (WT-CNO; *N* = 5 recordings). Initial inspection of the data revealed an increase in normalised spike count in the GlyT2-CNO groups, but not Veh and WT-CNO groups, shortly after application of CNO (**Figure 3c,d**). Additionally, when comparing the mean of the normalised post-drop cortical spike counts (5-minute bin width), we found a significant difference across conditions (ANOVA, *F* = 10.91, *p* = 2.82e-05). However, significant variability in spike counts were observed both within and between experimental groups. Therefore, to confirm that alterations in spike count observed across recordings were due to chemogenetic manipulation of GlyT2-positive cells, we used a multilevel model approach; this approach was warranted by the temporal correlations between repeated measures and clustering by animals. The model described the distribution of spike counts for each time bin with an inverse-Gamma distribution, parameterised by a shape parameter α and a scale parameter β. The model used as explanatory variables the experimental conditions, associated to pre- and post-drug administration periods, to find the α and β parameters that best explained the pre- and post-drug administration data. For each parameter of the gamma distribution (*α* and *β)* and each group (WT-CNO, Veh, GlyT2-CNO), the model had a coefficient *θ* associated to the pre-drug recording period and one for the post-drug administration period. The contrast between the two coefficients was used to measure the change in spike counts due to the experimental manipulation. To confirm the fidelity of the model, we compared the distribution of all experimentally observed spike counts together with the model fit (**Figure 3e**): the overlap between these distributions indicated that the model performed well in capturing the overall structure of the data.

Having confirmed the fidelity of the model, we then compared the contrasts between model coefficients associated with the pre- and post-drug administration periods for each experimental group. These contrasts reflect the effect of the experimental manipulation on spike count distribution. For each experimental group there are two contrasts, one associated with the shape parameter *α* and one with the scale parameter *β* of the distribution describing spike counts. Comparing contrasts and highest density intervals (hdi, 94%) for the parameters across groups revealed a specific effect in GlyT2-CNO mice on both *α* and *β* (**Figure 3f**). To gain a better intuition of the implications of these results, we used the posterior samples of *α* and *β* to derived samples for the mean and variance parameters of the inverse-Gamma distribution; the mean and variance offer an alternative parametrisation that characterise more intuitively the centre of mass and spread of spike counts across recordings for each condition, before and after drug application (see Materials and Methods). Thus, to compare results across condition, we computed the pre-minus post-drug mean and variance contrasts for each experimental condition, and the difference in contrasts between conditions, for each parameter. This analysis revealed that both mean and variance increased after drug application in the GlyT2-CNO group compared to the two control groups. (**Figure 3g**; mean hdi 0.05-0.46, 0.05 - 0.49 and variance hdi - 0.01 – 0.31, -0.02 – 0.31 for GlyT2-CNO versus WT-CNO and Veh respectively). It further revealed that in the WT-CNO group, CNO application alone has no detectable effect on network spiking activity (mean hdi -0.17– 0.21 and variance hdi -0.02 – 0.03 for WT-CNO versus Veh). Together our analyses confirm that chemogenetic inhibition of Golgi cells causes a global increase in spiking activity within the cerebellar cortex.

### Limited influence of GlyT2-positive cell downregulation on sensorimotor representation

Having established that chemogenetic downregulation of GlyT2-positive cells measurably altered cerebellar activity, we investigated how the perturbation affected sensorimotor representations of whisking. Golgi cell inhibition acts via the granule cell layer, but changes in the input-output transformation of granule cells will have knock on effects upon the activity of molecular interneurons and Purkinje cells. Golgi cells control spike output of granule cells through various mechanisms, including changes in gain, i.e., slope of the input-output function (Brickley et al., 1996; Chadderton et al., 2004; Hamann et al., 2002; Mitchell & Silver, 2003). By reducing Golgi cell inhibition, granule cells may become more sensitive to excitatory inputs, increasing their gain. We looked for changes in the sensitivity of cerebellar activity to whisker movement by comparing population tuning curves before and after CNO administration in GlyT2 mice. We observed no striking differences in the overall profile of these curves (**Figure 4a**). Post-CNO, population firing rates were marginally higher for some whisker positions, but tuning curves were essentially unchanged. This result suggests that chemogenetic downregulation of GlyT2-positive cells has no substantial effect on gain or overall representation of whisker position at the population level. In principle, altering Golgi inhibition could cause reorganisation of the granule cell population code by altering the requirements for granule cell integration of excitatory inputs from different mossy fibres (Marr, 1969). We therefore explored whether perturbed GlyT2-positive cell activity causes substantial reorganisation of whisking representation at the single cell level. To do this, we measured the entropy of individual tuning curves before and after CNO/vehicle administration. Entropy is a measure of how informative units are about, in this case, whisking position, with higher-entropy neurons being less informative. Following CNO administration, we observed no changes in cumulative entropy distribution in GlyT2-CNO recordings (**Figure 4b**). Interestingly, under control conditions (i.e. Veh and WT-CNO recordings), units tended to have higher entropy values after vehicle/CNO administration, indicating that tuning curves become less informative about whisking position over time. We further compared the distributions of entropy differences pre- and post-drug delivery (**Figure 4c**). Under control conditions, tuning curve entropies typically remained unchanged or increase over time (control density skewness = 2.80). CNO administration in GlyT2 mice causes additional reorganisation - some units decrease their entropy - leading to an even distribution of entropy changes after drug application (GlyT2-CNO density skewness = 0.21). The cumulative entropy distribution in GlyT2-CNO recordings therefore remains similar across time suggesting that the firing rate of some units becomes more sensitive to changes in whisking position following GlyT2-positive cell manipulation. These results also imply that recording conditions in the control condition are associated with small but progressive losses of representational fidelity. To confirm this proposal, we again measured the correlation between each of the first three population principal components (pc1-3) and whisker position for GlyT2-CNO and control recordings (**Figure 4d,e**). We found no evidence for large differences in correlation pre- and post-drug administration. However, there was a general trend for correlations to decrease after drug delivery, except for pc1 and pc2 (i.e. fast and slower representations of whisking) in the GlyT2-CNO condition, where correlations increased slightly. Our results suggest that the fidelity of sensorimotor representation is subject to a mild downward drift under control conditions in the present experimental setting, and that GlyT2-positive cell downregulation may slightly improve representation, offsetting this drift. But overall, we did not observe notable changes in the quality of the encoding of whisking behaviour by population activity. Together our results indicate that individual sensorimotor tuning of cerebellar neurons is only weakly influenced by chemogenetic perturbation of GlyT2-positive interneurons.

**Figure 4.**
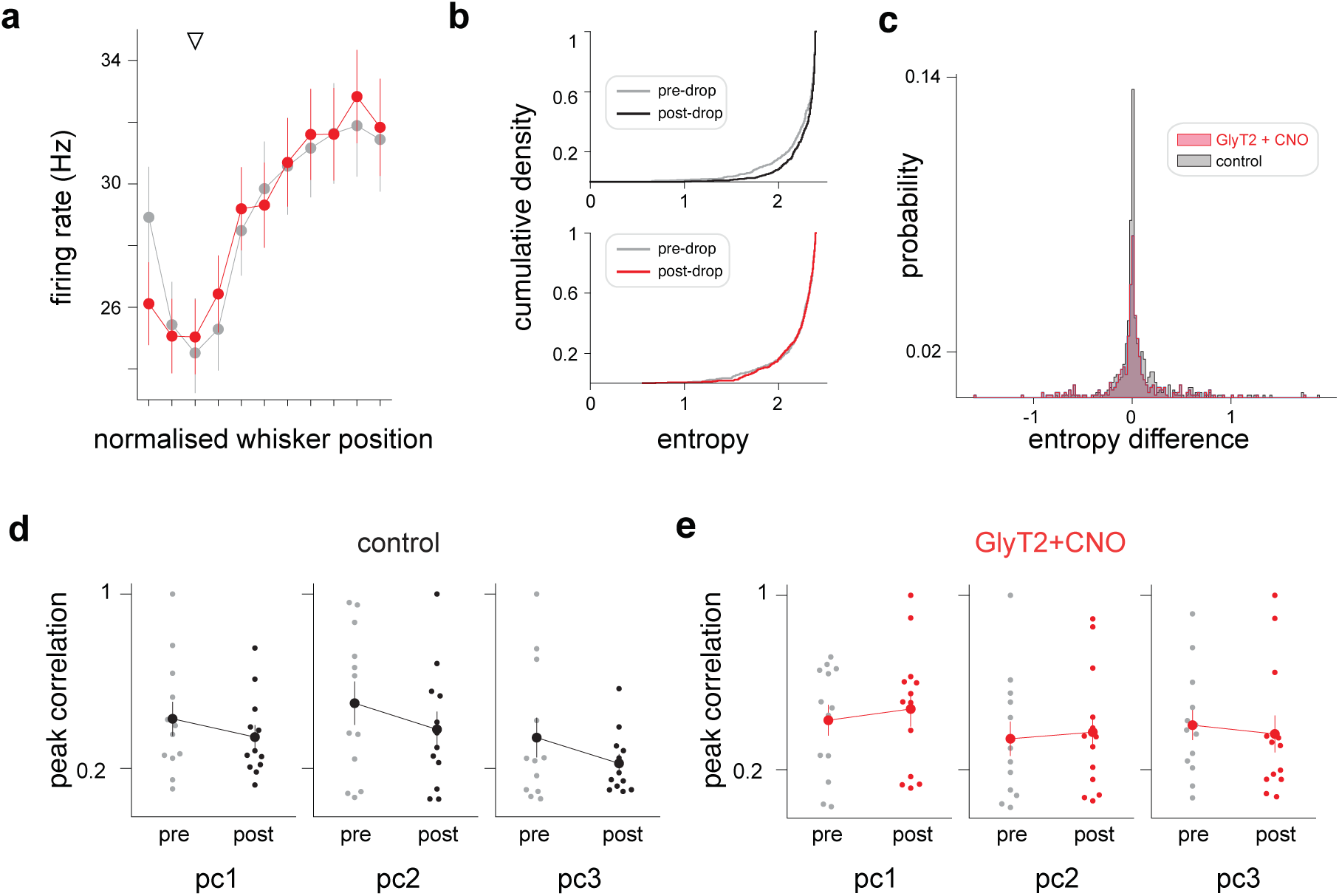
Weak influence of GlyT2-positive cell perturbation on cerebellar sensorimotor representations. **a.** Average tuning curves for GlyT2-CNO recordings before (grey) and after (red) topical administration of the drug. **b.** Entropy cumulative density functions for all units recorded in the GlyT2-CNO (top) and control (bottom) conditions. Under control conditions, but not GlyT2-CNO, unit entropy tends to increase after drug/vehicle delivery (post-drop), suggesting that tuning curves become less informative about whisking position over time. **c.** Probability mass function of unit entropy differences before and after CNO/vehicle administration (‘post-drop’ minus ‘pre-drop’ entropy). Under control conditions, tuning curve entropy tends to remain stable or increase over time (density skewness = 2.79). In the GlyT2-CNO recordings, a fraction units exhibit decreased entropy after drug application (density skewness = 0.21), suggesting that some units become more sensitive to changes in whisking position following GlyT2-positive cell inhibition. **d.** Comparison of peak pre- and post-drop correlation values between principal components 1-3 (pc1-3) and whisker position for control recordings. **e.** Comparison of peak pre- and post-drop correlation values between principal components 1-3 (pc1-3) and whisker position for GlyT2-CNO recordings.

### GlyT2-positive cell downregulation alters cerebellar dynamics and weakens coupling to whisking behaviour

We further explored the impact of GlyT2-positive cell perturbation by comparing trial averaged peri-event time histograms (PETH) and whisking activity from pre- and post-drug administration periods (see Materials and Methods). Because cerebellar activity can directly influence whisker movement (Proville et al., 2014), reduced GlyT2-positive cell inhibition could influence both cerebellar dynamics and produce measurable changes in whisking behaviour. We first considered cerebellar activity: reductions in Golgi cell inhibition could alter the temporal dynamics of population activity in several ways, acting via tonic and/or phasic inhibition (Brickley et al., 1996; Chadderton et al., 2004; Duguid et al., 2015), and/or affecting Golgi cell synchrony (van Welie et al., 2016). We therefore compared the patterns of population activity at movement onset before and after chemogenetic perturbation. Typically population activity peaks close to the onset of individual whisking bouts (**Figure 1b,c**). Therefore, for each recording, we computed the standard deviation of the distribution of peak unit firing times aligned to whisking onset, pre- and post-drug delivery (**Figure 5a**; see Materials and Methods). In the GlyT2-CNO mice, but not controls, pre- and post- drug standard deviations were significantly different (**Figure 5b**), with significantly lower variation following CNO administration (two-sided Wilcoxon signed-rank test, *T* = 12, *p* = 0.017). Thus, GlyT2- positive cell downregulation increased the temporal alignment of peak neuronal activity around whisking onset. These results suggest that decreased GlyT2-positive cell inhibition has an impact on the pacing of neuronal responses around the onset of whisking activity, and in particular narrows the time window within which units fire most strongly.

**Figure 5.**
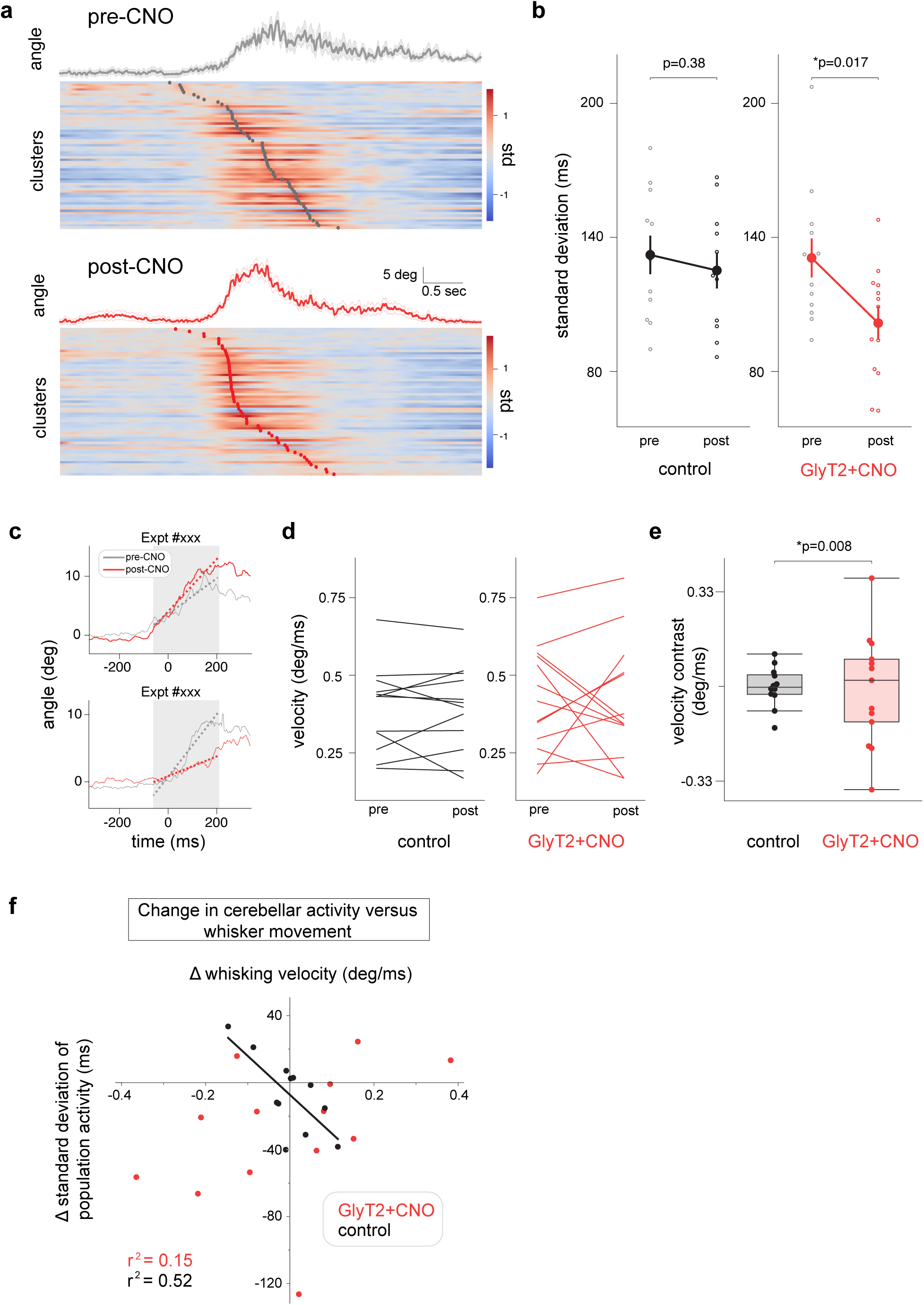
GlyT2-positive cell inhibition reduces temporal variability in neuronal populations and weakens behavioural coupling at movement onset. **a.** Peri-event time histograms aligned to onset of whisker movement for pre-(top) and post-drop (bottom) periods for a GlyT2-CNO recording. Grey and red dots indicate the absolute peak of neuronal activity centred around whisking onset for the pre- and post-drop period, respectively. Units ranked according to their peak firing rate. Temporal jitter peak activity decreases after CNO application. **b.** Temporal dispersion of neuronal population activity before and after drug/vehicle administration (pre- and post-drop) for control (*n* = 12) and GlyT2-CNO (*n* = 13) recordings. Each data point represents the standard deviation of the distribution of absolute peak times of neuronal activity for each recorded population. Reduction of GlyT2-positive cell inhibition decreases the temporal dispersion of neuronal activity during whisking initiation (two-sided Wilcoxon signed-rank test, *T* = 12, *p* = 0.017). **c.** Average whisking onsets for two representative GlyT2-CNO recordings before (grey) and after (red) drug administration. Dashed lines represent linear fits of initial whisker protraction. CNO delivery was associated with both increased (top) and decreased (bottom) slope of protraction. **d.** Change in slope of whisker protraction at movement onset before and after drug/vehicle administration (pre- and post-drop) for control and GlyT2-CNO recordings. **e.** Box plot showing differences in slope of whisker protraction at movement onset between pre- and post-drug/vehicle administration for control and GlyT2-CNO recordings. The standard deviation of the slope distribution in the GlyT2-CNO condition is higher than controls (Levene’s test *W* = 8.39, *p* = 0.008) suggesting that decreasing local GlyT2-positive cell inhibition produces variable changes in the dynamics of movement onset across recordings. **f.** Relationship between change in cerebellar population dynamics and whisker movement following drug/vehicle administration for all recordings. Under control conditions, measured changes in neural activity and movement are small in magnitude and well correlated (*r*^2^ = 0.52). In GlyT2-CNO recordings, relationships between cerebellar activity and movement are decoupled (*r*^2^ = 0.15).

Finally, we measured the effects of GlyT2-positive cell manipulation on whisking velocity at the onset of movement bouts (see Materials and Methods). For each recording, we used a linear model to approximate the slope of the average whisking bouts during its protraction phase (**Figure 5c**); the coefficient for each linear fit was used as a read out for velocity during the movement initiation. Interestingly, unlike control recordings, GlyT2-CNO mice showed changes in their whisking behaviour following drug administration. However, the sign of these changes was not consistent across animals, occurring in both increasing and decreasing directions (**Figure 5d**). Individual onsets of voluntary whisking thus became faster or slower after drug administration in GlyT2-CNO mice. The overall distribution of changes in whisker onset velocity (‘slope contrast’) was therefore significantly broader in the GlyT2-CNO mice compared to control (**Figure 5e**). Thus, GlyT2-positive cell downregulation produces measurable but divergent effects on local cerebellar dynamics and whole animal whisking behaviour. Local cerebellar dynamics were consistently compressed, leading to a more tightly locked population activity, whereas whisking velocity was either increased or reduced. These results suggest a breakdown in the link between cerebellar dynamics and behaviour following GlyT2-positive cell downregulation. We confirmed this by plotting changes in population activity and whisking velocity at movement onset for individual recordings (**Figure 5f**). In control recordings, differences in network activity and whisker movement before and after CNO/vehicle delivery are small in magnitude but importantly, are well correlated (*r*^2^ = 0.52). In GlyT2-CNO recordings, changes in cerebellar dynamics and movement become decoupled from one another, leading to break down in this correlation (*r*^2^ = 0.15). Together our results indicate GlyT2-positive cell inhibition regulates the temporal patterning of population activity and that disruption of this patterning at the local level can have diverse consequences for behaviour.

## DISCUSSION

Granule cell layer inhibitory interneurons are proposed to play important roles in cerebellar transformation (Albus, 1971; D’Angelo et al., 2013; Marr, 1969; Palacios et al., 2021), but we lack detailed information about their influence on circuit activity. Here we have tested the influence of Golgi and Lugaro cells on cerebellar population activity. Our GlyT2-positive cell perturbations caused mild changes in neural population activity without profoundly disrupting sensorimotor representations in the cerebellar cortex. Such an alteration is likely within normal physiological operating limits, and the changes we observe can thus be presumed to be ethologically relevant.

### Cerebellar cortical neurons heterogeneously encode whisking behaviour

Lateral cerebellar neurons accurately represent whisking behaviour. Individual units exhibit different tuning profiles to whisking position (**Figure 1d**), consistent with previous work (Bosman et al., 2010; Brown & Raman, 2018; Chen et al., 2016, 2017), enabling multidimensional representations of movement kinematics (Gurnani & Silver, 2021; Markanday et al., 2023). Our units are unlikely to be granule cells (Beau et al., 2024), but rather inhibitory interneurons and Purkinje cells. However, as observed in small virtual granule cell populations (Chen et al., 2017), ensemble activity displayed a near-monotonic dependence upon whisker set point, with increasing activity in the direction of both protraction and retraction (**Figure 1e**) in line with previous findings (Brown & Raman, 2018; Chen et al., 2016, 2017; Lanore et al., 2021; Proville et al., 2014).

### The cerebellar cortex uses a distributed code to represent whisking dynamics

We used principal component analysis to describe population activity with three independent variables. The first three pcs capture only a moderate amount of the total variance in the population, suggesting that cerebellar activity reflects additional behaviourally relevant variables (Chabrol et al., 2015; Ishikawa et al., 2015; Sobel et al., 1998; Wagner & Luo, 2020). Nevertheless, neuronal dynamics had structure that matched whisking dynamics (**Figure 1g**). Notably, pc1 dynamics consistently led (i.e. anticipated) whisking behaviour by ∼40ms whereas pc2 dynamics lagged behaviour by a few hundred milliseconds (**Figure 2**). The former is consistent with a forward model of planned/predicted movement, while the latter resembles a feedback signal. These results support a role for the cerebellum in controlling future behaviour, incorporating recent sensory feedback and further behaviourally relevant variables, in line with theoretical treatments of the cerebellum (Palacios et al., 2024). Whisker-related information, especially in pc1, was well distributed across units. Thus, most units contribute to some extent to its encoding.

### Decreasing GlyT2-positive cell inhibition does not saturate population activity

Next, we examined how altering GlyT2-positive cell activity affected neuronal representations in the cerebellar cortex. This was achieved by activating inhibitory receptors (hM4Di) on Golgi and Lugaro cells, through application of the agonist CNO directly at the surface of the recording site. This approach afforded within-minute temporal precision to the manipulation. hM4Di receptors inhibit neuronal activity via moderate hyperpolarisation and reduced presynaptic release of neurotransmitters (Roth, 2016). Topical application of CNO onto the cerebellar cortex produced an increased spread of the population spike count distribution, with higher spike counts becoming more likely across recordings (**Figure 3c,d**). However, manipulation of GlyT2-positive cell inhibition did not lead to excitatory saturation of network dynamics, i.e., a drastic and indiscriminate increase in neural activity. The absence of strong saturation of network activity may be because granule cell excitability is partly controlled via non-vesicular sources of GABA (Lee et al., 2010; Rossi et al., 2003; Wall & Usowicz, 1997). In addition, Lugaro cells target inhibitory neurons, here potentially producing compensatory upregulation of molecular layer interneuron activity (Eyre & Nusser, 2016). The tuning of individual neurons was also largely unchanged (**Figure 4a**). Theoretical and *in vitro* evidence indicates that the gain of granule cell input/output transformation may be under the control of local inhibitory processes. As such, reduced inhibition is predicted to increase cortical sensitivity to incoming synaptic input. Following our perturbation, we saw little evidence of increased sensitivity to features of whisker movement (**Figure 4a**), although we did observe a small fraction of neurons that exhibited reduced entropy following GlyT2-positive cell inhibition (**Figure 4c**). These neurons may represent a subpopulation that receive subthreshold whisker-related synaptic input. Here, small decreases in synaptic inhibition may result in the generation of tuned action potential output (Chadderton et al., 2004; Chen et al., 2017). However, our results suggest that dynamic GlyT2-positive cell inhibition may not be necessary for maintaining excitation at the level required by the network to perform its computations (Albus, 1971; Marr, 1969). Manipulation of GlyT2-positive cell inhibition instead produces more fine-grained effects on cerebellar cortical dynamics.

### Decreased GlyT2-positive cell inhibition increases the temporal alignment of neuronal activation during movement onset

Decreased GlyT2-positive cell inhibition was associated with increased temporal alignment or synchronisation within neuronal populations around whisking onset. A familiar downstream consequence of feedforward inhibition in many brain regions, and at the molecular layer interneuron-Purkinje cell synapse, is increased temporal precision of neuronal dynamics, as inhibition normally trails excitation and sharpens the window of input integration (Gabernet et al., 2005; Mittmann et al., 2005; Wehr & Zador, 2003). Here, notably, Golgi cells play the opposite role, temporally dispersing or desynchronising neural activity (Vervaeke et al., 2010). Although Golgi cells provide feedforward inhibition to the granule cell layer, evoked inhibition has been previously shown to precede or be concomitant with mossy fibre excitation, under which situations it may reduce the temporal precision of early granule cell responses (Duguid et al., 2015), and decorrelate multisensory representations (Fleming et al., 2024). Our observations are consistent with such a mode of operation, as decreasing GlyT2-positive cell inhibition leads to an increased temporal alignment of the overall network activity during movement initiation (Vervaeke et al., 2010). Changes in Golgi cell activity can modulate the amplitude of tonic inhibitory conductances (Crowley et al., 2009). Reduced Golgi cell activity may therefore directly reduce the tonic inhibitory conductance, making granule cells more excitable and more likely to fire rapidly to a burst of incoming mossy fibre synaptic inputs. Such control by Golgi cells of the timing of network activity can support the specific cerebellar role in integrating information and learning contexts (Fleming et al., 2024). Indeed, distinct temporal patterning of granule cell responses to mossy fibre input is proposed to be important for how the cerebellar network integrates and distinguishes parallel sensory and motor pathways (Chabrol et al., 2015).

### Decreasing GlyT2-positive cell inhibition decreases the stability of whisking behaviour

Finally, we found that reduced GlyT2-positive cell inhibition led to increased variability in whisking velocity at movement onset. The specific change in movement velocity varied amongst recordings: in some cases, the velocity of whisking increased while in other cases it decreased. We suggest that location of the site of perturbation, varying from mouse to mouse, is likely to have influenced the valence of the change in whisking behaviour (Apps et al., 2018). Importantly, the link between cerebellar cortical dynamics and whisker position is decoupled after GlyT2-positive cell perturbation (**Figure 5f**). This result highlights the importance of an intact cerebellum for precise whisking control, which is subject to the coordination with other motor and sensory systems (Deschenes et al., 2012; Kurnikova et al., 2017; McElvain et al., 2018; Romano et al., 2020). Increased variability in motor initiation may be linked with the possible contribution of Golgi cell inhibition to representations of multiple streams of information and behaviours – bound together at the level of the cerebellar cortex – improving accuracy as specific behaviours are performed.

### Conclusions

Granule cell layer inhibition itself is under the control of many mechanisms, including feedback loops encompassing the cortex (Nietz et al., 2017; Witter et al., 2016), the cerebellar nuclei (Ankri et al., 2015), as well as neuromodulatory systems (Dieudonne & Dumoulin, 2000; Fleming et al., 2024; Fore et al., 2020). Thus, granule cell layer inhibitory interneurons sit in a key position to flexibly control cerebellar computations, based on the dynamic behavioural context, which may be required to focus on different aspects of behaviour at different times (Palacios et al., 2021). To fully understand the pervasive involvement of the cerebellum in behavioural control, it is critical to elucidate how different mechanisms contribute to fine tune Golgi and Lugaro cell activity in a context-dependent manner.

Our work shows how altering GlyT2-positive cell inhibition changes the response of the network to its input. By controlling the timing of network activity, GlyT2-positive interneurons can support cerebellar roles in integrating information and learning contexts (Fleming et al., 2024). Distinct temporal patterning of granule cell responses to mossy fibre input may enable the cerebellar cortex to integrate and distinguish parallel sensory and motor pathways, which in turn may be relevant for pattern separation (Chabrol et al., 2015). Cerebellar computations have also been described as state estimation processes (Doya, 1999; Miall et al., 1993; Palacios et al., 2024; Paulin, 1993) in which granule cell layer inhibition acts as a mechanism to fine tune the integration of information from different mossy fiber pathways (Palacios et al., 2021). In this scheme, temporal changes in cerebellar dynamics may result from tuned integration of whisking-related information with other, temporally heterogeneous streams of information.

## AUTHOR CONTRIBUTIONS

ERP, CH and PC designed the experiments. All experiments were performed by ERP. Data was analysed and interpreted by ERP, CH and PC. The authors declare no competing interests.

## ACKNOWLEDGEMENTS

This work was supported by a Wellcome Trust Neural Dynamics PhD studentship to ERP, Leverhulme Research Fellowship (RF-2021-533) to CH and Wellcome Trust Investigator Award (209453/Z/17/Z) to PC. The schematic images in Figure 1 were generated and provided by Elisabeth Meyer whom we also thank for comments on the manuscript.

The authors declare no competing interests.

