## Supplementary material for "GlyT2-positive interneurons regulate timing and variability of information transfer in a cerebellar-behavioural loop": Table 1

| Recording | Animal | Strain | Drug | Whisking | # cortical units | Time CNO/PBS drop (min) |
|---|---|---|---|---|---|---|
| 1 | 1 | GlyT2 | CNO | poor | 18 | 10 |
| 2 | 1 | GlyT2 | CNO | good | 5 (4) | 10 |
| 3 | 2 | GlyT2 | CNO | good | 8 (7) | 10 |
| 4 | 2 | GlyT2 | CNO | good | 12 (11) | 10 |
| 5 | 3 | GlyT2 | CNO | good | 10 | 10 |
| 6 | 3 | GlyT2 | CNO | good | 10 (6) | 10 |
| 7 | 4 | GlyT2 | CNO | good | 8 | 20 |
| 8 | 5 | GlyT2 | CNO | absent | 27 | 20 |
| 9 | 6 | GlyT2 | CNO | good | 61 | 20 |
| 10 | 7 | GlyT2 | CNO | good | 76 | 20 |
| 11 | 7 | GlyT2 | CNO | good | 24 | 20 |
| 12 | 8 | GlyT2 | CNO | absent | 5 | 20 |
| 13 | 8 | GlyT2 | CNO | poor | 30 | 20 |
| 14 | 9 | GlyT2 | CNO | good | 58 | 20 |
| 15 | 9 | GlyT2 | CNO | good | 23 | 20 |
| 16 | 10 | GlyT2 | CNO | good | 46 | 20 |
| 17 | 10 | GlyT2 | CNO | good | 49 | 20 |
| 18 | 11 | GlyT2 | CNO | poor | 9 | 20 |
| 19 | 11 | GlyT2 | CNO | poor | 29 | 20 |
| 20 | 12 | WT | CNO | good | 96 (88) | 10 |
| 21 | 13 | WT | CNO | good | 28 | 20 |
| 22 | 13 | WT | CNO | good | 31 | 20 |
| 23 | 14 | WT | CNO | good | 14 | 20 |
| 24 | 14 | WT | CNO | good | 55 (54) | 20 |
| 25 | 4 | GlyT2 | PBS | good | 8 | 20 |
| 26 | 5 | GlyT2 | PBS | absent | 23 | 20 |
| 27 | 6 | GlyT2 | PBS | good | 31 | 20 |
| 28 | 15 | WT | PBS | good | 88 | 20 |
| 29 | 15 | WT | PBS | poor | 38 | 20 |
| 30 | 16 | WT | PBS | good | 61 | 20 |
| 31 | 16 | WT | PBS | good | 28 | 20 |
| 32 | 17 | WT | PBS | good | 52 | 20 |
| 33 | 17 | WT | PBS | good | 25 | 20 |

**Supplementary Table 1**: Neuronal and behavioural data. Each animal underwent two recordings (except animal 12). Each recording was performed on either GlyT2-Cre or C56BL6 (wild-type; WT) mice; GlyT2 mice expressed Cre-recombinase selectively in Golgi cells in the cerebellar cortex, and therefore only in these mice CNO could selectively decrease Golgi cell inhibition. The three experimental conditions are GlyT2+CNO, in which CNO was used on GlyT2-Cre mice, WT+CNO in which CNO was used on WT mice, and vehicle, in which phosphate-buffered saline (PBS) was used on either injected GlyT2-Cre mice (3 recordings) or WT mice (6 recordings); the WT+CNO and vehicle conditions were pooled into one control condition, after assessing for the specific effect of our manipulation on total cerebellar cortical spike counts using the statistical model described in equations 1-4 (see *Materials & Methods*). For this statistical analysis of spike counts, we used all units from all recordings ($n$ = 1086 units, $N$ = 33 recordings). In all other analyses, instead, we excluded recordings with absent/poor whisking activity ($N$ = 25 after exclusion), as the analyses required concomitant behavioural and neuronal information; for the same reason, we additionally excluded a small number of units ($n$ = 16, remaining number of units in parenthesis) whose activity was absent or too sparse during whisking periods in order to compute the peri-event time histogram. Poor whisking behaviour was defined as a flat trial-averaged whisker position trace for either or both pre- and post- drop periods. Control mice only were used for analysis included in Figures 1 and 2 ($n$ = 508, $N$ = 12 recordings after exclusion).
